# The EphB2-MYC Axis is a Major Determinant of Barrett’s Pathobiology and a Therapeutic Vulnerability in Esophageal Cancer

**DOI:** 10.1101/2021.05.13.444044

**Authors:** Srividya Venkitachalam, Deepak Babu, Durgadevi Ravillah, Ramachandra M. Katabathula, Peronne Joseph, Salendra Singh, Bhavatharini Udhayakumar, Yanling Miao, Omar Martinez-Uribe, Joyce A. Hogue, Adam M. Kresak, Dawn Dawson, Thomas LaFramboise, Joseph E. Willis, Amitabh Chak, Katherine S. Garman, Andrew E. Blum, Vinay Varadan, Kishore Guda

## Abstract

Esophageal adenocarcinoma (EAC), a highly aggressive cancer with limited therapeutic options, often arises in the backdrop of a molecularly-complex esophageal metaplasia disorder, Barrett’s Esophagus (BE). Using transcriptomics and systems biology analyses of treatment-naïve malignant/pre-malignant biopsy tissues, we found Eph receptor B2 (EphB2) tyrosine kinase signaling to be frequently hyperactivated during early stages of EAC development, and across the BE-EAC continuum. Functional studies revealed EphB2 to be an upstream post-translational regulator of c-MYC activity and as a key molecular dependency in BE/EAC. Single-cell transcriptomics in a porcine esophageal 3D spheroid model showed enhanced EphB2 and MYC activity to be significantly associated with BE-like cell fate. shRNA-based knockdown of EphB2 or small molecule inhibitors of MEK, that modulate MYC protein stability, proved effective in suppressing EAC tumor growth *in vivo*. These findings point to EphB2-MYC axis as an early promoter of EAC and a novel therapeutic vulnerability in this increasingly-prevalent esophageal malignancy.

**STATEMENT OF SIGNIFICANCE:** We identify EphB2 signaling as a potential master regulator and early promoter of esophageal adenocarcinoma, and the proto-oncogene MYC as a key downstream effector of EphB2 function. Targeting the EphB2-MYC axis could be a promising therapeutic strategy for these often refractory and lethal EAC tumors.

## INTRODUCTION

Barrett’s esophagus (BE) is a clinical condition wherein the esophageal squamous epithelia (SQ) is replaced by intestinal-type columnar epithelia, often in response to mucosal injury from chronic gastrointestinal reflux disease (GERD) (1). Importantly, patients with BE are at an increased risk for developing esophageal adenocarcinoma (EAC), a lethal malignancy with increasing incidence rates even in younger populations within the U.S. (2,3). Moreover, the vast majority of EAC cases are refractory to standard of care treatments (4), and targeted therapies are virtually non-existent (4,5), resulting in a dismal 5-year survival (3,5). Despite the significance of BE and associated neoplasia, advancements in the clinical management of this disease have been stymied due to incomplete understanding of unifying mechanisms driving the onset and progression of BE pathogenesis.

Herein, we performed integrative genome-scale analyses of transcriptomic profiles derived from treatment-naïve pre-malignant and malignant biopsy tissues, followed by pharmacogenetic studies, identifying EphB2-MYC as a key signaling axis in BE pathogenesis, with potential therapeutic implications.

## RESULTS

### EphB2 signaling is hyperactivated in the vast majority of BEs and EACs

In order to identify early drivers of BE/EAC, we employed a systems biology framework, InFlo (6,7), to analyze RNA sequencing (RNASeq) data derived from a discovery cohort (8) consisting of 49 pre-treatment EACs, 18 Non-Dysplastic Stable Barrett’s Metaplasia (NDSBM) and 11 normal esophageal squamous tissues (nSQ) (**Supplementary Table S1**). Besides known alterations in BE-associated pathways (See Methods), we found Eph Receptor B2 (EphB2) Tyrosine Kinase signaling to be hyperactivated in 88% of EAC and 100% of NDSBM samples as compared to nSQ (**Fig. 1A**). Consistent with this, we found EphB2 signaling components including, EphB ligands (*EFNB1* and *EFNB2*) and *EPHB2* receptor as being significantly induced in NDSBM and EAC lesions, in comparison to nSQ (**Fig. 1B**).

**Figure 1.**
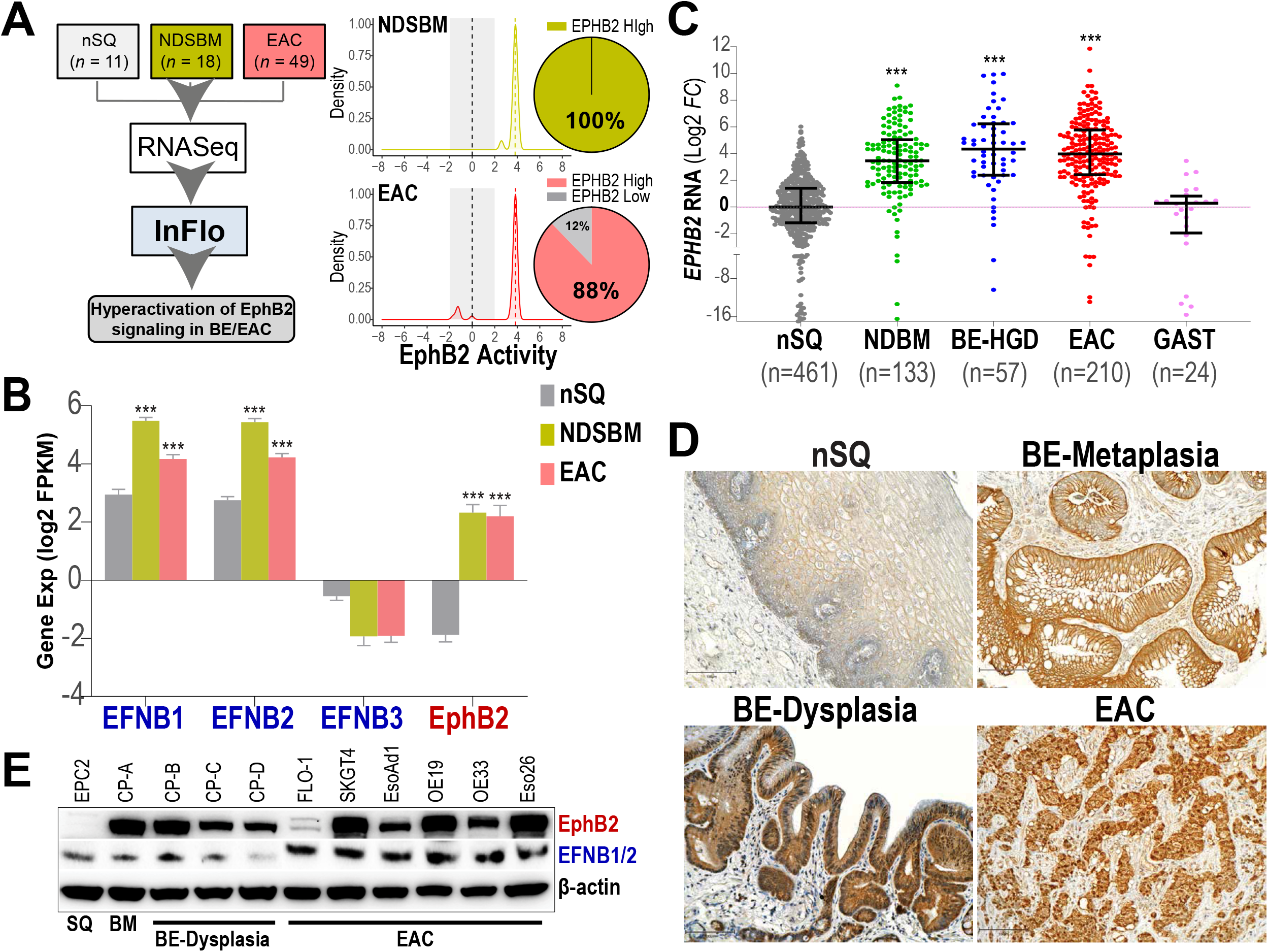
EphB2 signaling is hyperactivated in a majority of BE and EACs. **A**, Analysis of RNASeq data derived from normal esophageal squamous (nSQ), Non-Dysplastic Stable Barrett’s Metaplasia (NDSBM), and Esophageal Adenocarcinoma (EAC) using the InFlo systems biology framework identified hyperactivation of EphB2 signaling in the vast majority of NDSBM and EAC when compared to nSQ tissues. Plotted are the distributions of InFlo-derived EphB2 signaling activity levels for NDSBM (yellow) and EAC (light red) tissue samples. InFlo-derived EphB2 activity levels are plotted along the X-axis, with activities associated with nSQ (- 2 to +2) denoted in grey, and hyperactivation (EphB2 High) corresponding to EphB2 Activity levels > 2. Pie-charts indicate the percentage of NDSBM (*N* = 18) and EAC (*N* = 49) samples exhibiting EphB2 hyperactivation (EphB2 High). **B**, Bar-graphs depict gene expression levels of *EPHB2* receptor and its associated ligands (*EFNB1, EFNB2* and *EFNB3*) in nSQ, NDSBM and EAC primary tissues included in the discovery RNASeq cohort. Y-axis shows the expression levels in log2FPKM units within each tissue-type detailing the median ± interquartile range of the expression. *** (*P* < 0.0005) indicates statistical significance of differences in gene expression between NDSBM/EAC versus nSQ, estimated using a Student’s t-test assuming unequal variances. **C**, qPCR analyses of *EPHB2* gene expression in an independent cohort of treatment-naïve primary biopsy tissues consisting of EAC, non-dysplastic Barrett’s metaplasia with no longitudinal follow-up (NDBM), Barrett’s esophagus with high grade dysplasia (BE-HGD), as compared to nSQ. Normal gastric (GAST) samples were used as additional controls. Y-axis shows the fold-changes in *EPHB2* gene expression across samples, relative to the median expression in SQ cases. Horizontal lines within each tissue-type indicate the median ± interquartile range of fold-changes in expression. *** (*P* < 0.0005) indicates statistical significance of differences in gene expression between NDBM/BE-HGD/EAC versus nSQ, estimated using a Student’s t-test assuming unequal variances. **D**, Representative Immunohistochemistry images in FFPE tissue sections demonstrating marked increase in EphB2 protein expression in NDBM, BE-HGD and EAC lesions, as compared to nSQ (Scale bar, 100µm). **E**, Western blot images depicting protein levels of EphB2 and its ligands (EFNB1/B2), with β-actin as a loading control, across distinct EAC (FLO-1, SKGT4, EsoAd1, OE19, OE33, Eso26), BE-Dysplasia (CP-B, CP-C, CP-D), BM (CP-A) and normal SQ (EPC2) cell lines, demonstrating marked increase in EphB2 protein expression in near-all BE/EAC cell lines as compared to normal SQ. Of note, while FLO-1, SKGT4, EsoAd1 and OE33 lines were derived from esophageal adenocarcinomas in the distal esophagus, OE19 and Eso26 were derived from gastro-esophageal junctional cancers.

Eph Receptor Tyrosine Kinases impact diverse cellular processes across tissue types (9), with potential therapeutic and biomarker potential (10,11). Nonetheless, their significance in the context of BE neoplasia remain poorly understood. We accordingly prioritized EphB2 signaling, a previously uncharacterized pathway in BE-EAC, for further studies. To first assess the generality of our findings, we performed qPCR-based expression assessments of *EPHB2* in a large independent cohort (**Supplementary Table S2**) of treatment-naïve biopsy tissues (*N*=885), including, EAC, non-dysplastic Barrett’s metaplasia (NDBM), BM with high-grade dysplasia (BE-HGD), normal gastric (GAST) and normal esophageal squamous (nSQ) samples. Indeed, we found significant (*P<0*.*0005*) overexpression of *EPHB2* in NDBE as well as in progressive disease-stages (HGD and EAC), compared to normal SQ tissues (**Fig. 1C**). Consistent with this, immunohistochemical analyses in primary, treatment-naïve, tissue sections confirmed the marked induction of EphB2 protein in NDBM, BE-HGD and EAC as compared to nSQ (**Fig. 1D**). Furthermore, Immunoblot analysis revealed markedly higher EphB2 protein expression in nearly all patient-derived BE/EAC cell line models, as compared to a normal esophageal SQ cell line (**Fig. 1E**); whereas EFNB1/B2 ligands were expressed in all cell lines (**Fig. 1E**). Of note, these patient-derived EAC cell lines include the distal esophagus adenocarcinomas (EAC: FLO-1, SKGT4, EsoAd1, OE33) and also gastro-esophageal junctional cancers (GEJ: OE19, Eso26).

### BE/EAC cells exhibit significant growth dependency on EphB2 signaling

We next explored the functional consequences of EphB2 signaling in well-characterized, patient-derived EAC cell line models. Transient siRNA-based knockdown of *EPHB2* (See Methods) impeded both long-term colony growth as well as short-term cell growth of EAC cells (**Fig. 2A**). Reprising the *in vitro* growth dependencies, shRNA-based stable knockdown of *EPHB2* led to a marked reduction in the growth of EAC cells as tumor xenografts *in vivo* (**Fig. 2B**). To assess whether these dependencies are retained even at earlier stages of EAC development, we performed *EPHB2* siRNA knockdown studies in pre-malignant BE cell line models. Similar to our observations in EACs, knockdown of *EPHB2* suppressed cell viability of metaplastic as well as dysplastic BE cells (**Supplementary Fig. S1**). These findings, taken together with EphB2 signaling assessments in primary tumor cohorts (**Fig. 1**), support EphB2 as contributing to and sustaining BE/EAC cell growth from the earliest metaplastic stage, through the later dysplastic and neoplastic stages of Barrett’s esophageal adenocarcinoma.

**Figure 2.**
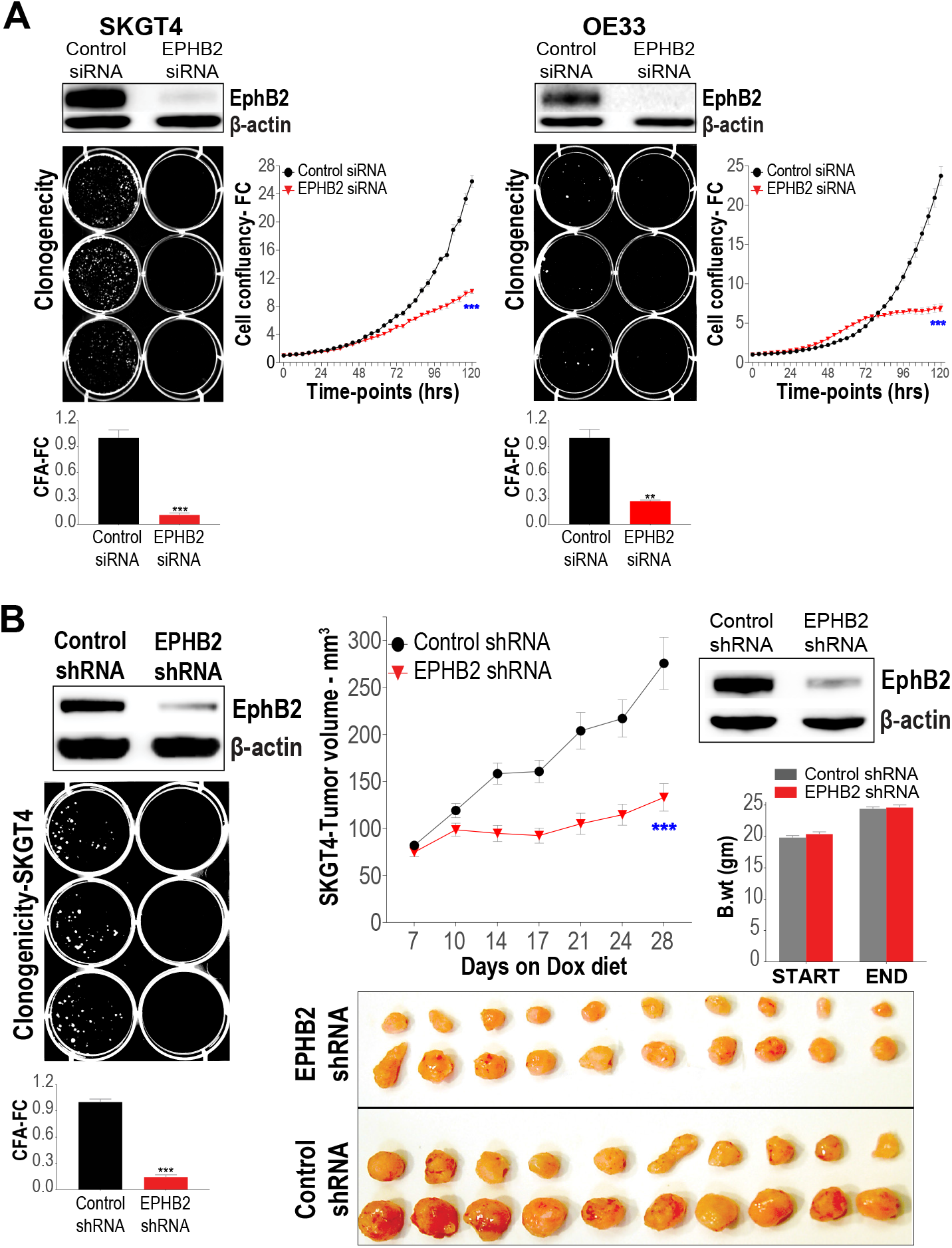
EAC cell growth is dependent on EphB2 signaling. **A**, Assessment of long-term EAC cell colony growth (left) and short-term IncuCyte-based growth assessments (right) upon siRNA-based *EPHB2* knockdown in representative EAC cell lines (SKGT4, OE33). On top are Western blot images depicting EphB2 protein expression 48hrs following treatment with either siRNA targeting *EPHB2* (*EPHB2* siRNA) or non-targeting control (Control siRNA). Shown on the left are representative images and bar-graphs of EAC cell colony numbers 10-14 days following siRNA treatment, expressed as a fraction of colony counts (CFA-FC) observed in respective cells treated with Control siRNA. Also shown are IncuCyte-based growth assessments over time (X-axis) of EAC cells depicting average fold-change in cell confluency values normalized to time zero (Cell confluency-FC), following either *EPHB2* siRNA or non-targeting Control siRNA treatment. All data are plotted as mean ± SEM, obtained from three independent experiments. ** (*P* < 0.005) and *** (*P* < 0.0005) indicate significant differences in growth assessments at the final timepoint estimated using a Student’s t-test assuming unequal variances. **B**, Assessment of *in vitro* clonogenicity and *in vivo* xenograft growth of SKGT4 (EAC) cells expressing either Doxycycline-inducible *EPHB2*-targeting or Control shRNA constructs. On the left are shown representative images and bar-graphs of colony growth in SKGT4 cells harboring either *EPHB2* shRNA or Control shRNA after 14 days of Doxycycline (Dox) treatment. Western blot images depict EphB2 protein expression 48hrs following Dox-treatment. Shown on the right are tumor growth kinetics of SKGT4 cells, expressing either Control or *EPHB2* shRNA, in immune-deficient mice fed with Dox-diet (See Methods). Y-axis depicts tumor volume in mm^3^ over time (X-axis) in respective SKGT4 xenografts. Data are plotted as mean ± SEM estimated using at least 20 established xenograft tumors at day zero in respective arms. *** (*P* < 0.0005) indicates significant differences in tumor volume at the final time-point between respective groups, estimated using a Student’s t-test assuming unequal variances. Western blot images depict EphB2 protein expression in representative tumor xenografts harvested after one week following Dox-treatment. Also shown are assessments of body weight of mice at the beginning and at the end of the study showing no marked differences in body weights of mice in either arm over the study duration. At the bottom are shown photographic images of harvested tumors from mice at the final time-point in both the *EPHB2* shRNA and Control shRNA arms.

### EphB2 signaling regulates MYC and its associated transcriptional program in BE/EAC

We next assessed for likely effector pathways of EphB2 signaling that potentially mediate EphB2 phenotypic effects in BE/EAC (**Fig. 2**). Accordingly, we performed RNASeq profiling and InFlo-based integrative analysis in representative EAC and BE cell lines, upon siRNA knockdown of *EPHB2* (See Methods). Besides previously-implicated pathways in BE/EAC such as NF-κB/RelA and Wnt/β-catenin (**Supplementary Fig. S2**), we in particular found the c-MYC proto-oncogene and associated transcriptional program to be significantly and positively regulated by EphB2 (**Fig. 3A**), uncovering a novel link between EphB2 and MYC. Interestingly, this regulation of MYC by EphB2 appeared to be at the post-translational level, as *EPHB2* knockdown resulted in marked reduction in the levels of MYC protein but not RNA across representative EAC and BE cell line models (**Fig. 3A**). To further test this association between EphB2 and MYC in primary lesions, we performed VIPER-based assessments of MYC transcriptional activity in our RNASeq dataset derived from BE and EAC tissue biopsies. Indeed, we observed a significantly higher MYC activity in EphB2-hyperactivated BE (*P*<0.0005) and EAC (*P*<0.005) samples, as compared to nSQ, despite no significant differences in *MYC* RNA expression between the tissue samples (**Fig. 3B**). These findings strongly point to EphB2 as a key post-translational regulator of MYC activity, and uncover a EphB2-MYC regulatory axis in the BE-EAC disease context.

**Figure 3.**
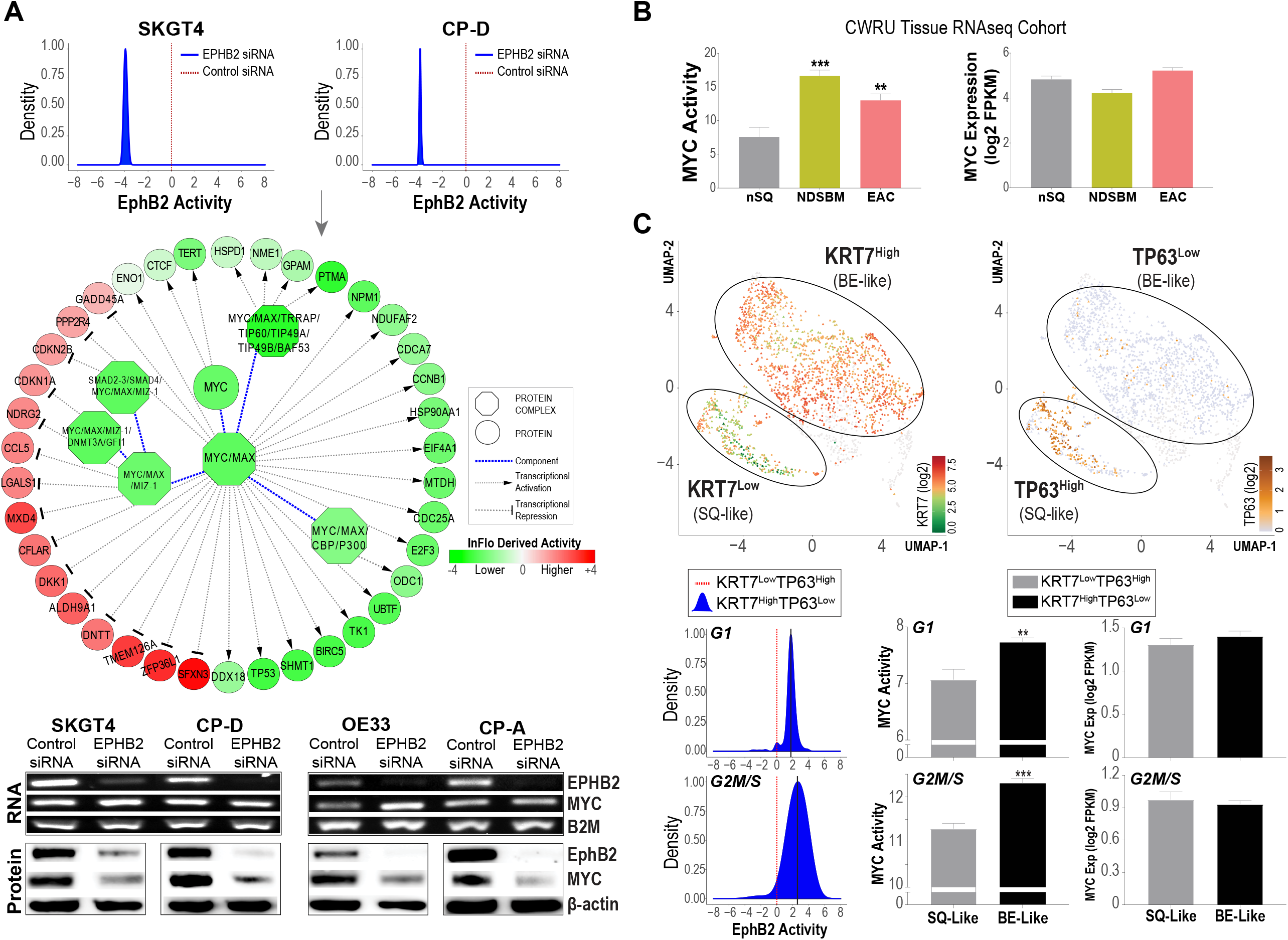
EphB2 signaling modulates MYC transcriptional activity in BE/EAC. **A**, InFlo-based analysis of whole transcriptome RNASeq performed on EAC (SKGT4) and dysplastic-BE (CP-D) cells following siRNA-based knockdown of *EPHB2* confirmed significant downregulation of EphB2 signaling upon treatment with *EPHB2* siRNA when compared to non-targeting Control siRNA (top). InFlo revealed a marked reduction in the activities of MYC protein complexes and their associated transcriptional targets (middle) upon *EPHB2* KD in both EAC/BE cells. Note the reduction (green) in activity of genes known to be transcriptionally activated by MYC, with a concomitant increase (red) in activity of genes known to be transcriptionally repressed by MYC. Shown at the bottom are images of qPCR products on agarose gel depicting *EPHB2* and *MYC* RNA expression, and Western blot images depicting EphB2 and MYC protein levels upon *EPHB2* and Control siRNA treatments in both SKGT4 and CP-D cells. Note the marked reduction in the levels of MYC protein but not RNA expression upon *EPHB2* knockdown even in OE33 (EAC) and CP-A (metaplastic BE) cells. *B2M* and β-actin were used as loading controls for qPCR and Western blots, respectively. **B**, Bar-graphs depicting MYC transcriptional activity assessments (top) using the VIPER framework (See Methods) and MYC gene expression levels (bottom) in nSQ, NDSBM and EAC primary tissues included in our discovery RNASeq cohort (See Fig. 1A). **C**, UMAP plots (top) of single-cell RNA sequencing (sc RNASeq) data generated from the porcine ESMG 3D culture model showing two distinct cellular clusters. Individual cells are colored according to their KRT7 (left) and TP63 (right) gene expression, thus classifying them into an SQ-like (KRT7^Low^ & TP63^High^) and BE-like (KRT7^High^ & TP63^Low^) population. The cell cycle phase of individual cells are denoted as circles (G1 phase) or triangles (G2M or S phase). The density plots (blue) denote distributions of InFlo-derived EphB2 signaling activities (X-axis) in BE-like KRT7^High^ & TP63^Low^ cells belonging to either the G1 (top) or G2M/S (bottom) phases, as compared to respective SQ-like manifolds (red dotted line) defined by KRT7^Low^ & TP63^High^ cells of the same cell phase. Bar graphs denote MYC activity and MYC expression level comparisons between BE-Like versus SQ-Like cells within the G1 (left) and G2M/S (right) populations, respectively. All data are plotted as mean ± SEM, with ** (*P* < 0.005) and *** (*P* < 0.0005) indicating statistical significance of differences estimated using a Student’s t-test assuming unequal variances.

### Activation of EphB2-MYC axis is an early event in BE pathogenesis

Our findings of hyperactivated EphB2 signaling and MYC activity even in early metaplastic stages of the BE-EAC disease continuum (**Figs. 1, 3B** and **Supplementary Fig. S1**) provocatively suggested that EphB2-MYC axis may be an early mediator of BE pathogenesis. Since Eph signaling has previously been implicated in abnormal tissue injury and repair (12), processes that are highly relevant in the etiology of BE (13), we evaluated the EphB2-MYC axis in esophageal submucosal glands (ESMGs), a cluster of progenitor cells implicated in esophageal repair as well as BE development (13). In particular, given that abnormal repair following injury can drive ESMGs to differentiate into BE-like columnar rather than SQ-like epithelial cells (14), we employed a porcine-derived 3D culture model of ESMGs to first determine whether EphB2 signaling is induced in early stages of transformative BE-like cell fate. We performed single-cell RNA sequencing (scRNASeq) in this 3D culture model, and subsequently employed a modified InFlo framework optimized for assessing differential signaling network activities in single-cells (See Methods). Strikingly, these analyses revealed significantly higher EphB2 activity in BE-like (*KRT7*^High^*TP63*^Low^), as compared to SQ-like (*KRT7*^Low^*TP63*^High^) spheroids, independent of cell-cycle phase (**Fig. 3C**). Concomitant with this, we again observed significantly higher MYC transcriptional activity (*P*<0.005) with no difference in *MYC* RNA expression levels *per se* in BE-like versus SQ-like cells, irrespective of cell-cycle phase (**Fig. 3C**). These findings strongly suggest that EphB2-MYC axis is an early modulator of cell fate in the etiology of BE.

### Small molecule inhibitors of MEK1 suppress MYC activity and impede EAC tumor growth in vivo

Our studies thus far identify MYC as a key downstream effector of EphB2 signaling in BE-EAC development, provocatively suggesting the EphB2-MYC axis as a potential therapeutic vulnerability in this disease. Given that there are no clinically-approved inhibitors specifically against EphB2 or c-MYC, we explored for surrogate druggable-targets involved in upstream-regulation of MYC activity. The MAPK signaling cascade emerged as a potential candidate given its known role in post-translationally regulating MYC stability and activity via ERK-mediated (Ser 62) phosphorylation of MYC (15,16), and our finding of EphB2 modulating the activity of multiple components of the MAPK signaling cascade in BE/EAC cell lines (**Supplementary Fig. S2**). We accordingly evaluated the *in vivo* anti-tumor efficacy of a clinically-approved potent MEK1 inhibitor (Cobimetinib) in pre-clinical tumor xenograft models. Sub-cutaneous tumor xenografts were established from the SKGT4 EAC cell line in immune-deficient (NCr-Foxn1^nu^ nude) mice. Once tumors reached a minimum size of 50-60mm^3^, respective mice were randomized to either treatment with Cobimetinib or vehicle alone and followed longitudinally (see Methods). Strikingly, Cobimetinib treatment resulted in strong suppression and/or marked regression of both EAC-derived xenograft tumors *in vivo*, without any detectable adverse effects on mice or on their body weights throughout the study duration (**Fig. 4**). Consistent with this, we found a marked reduction in the pharmacodynamic marker, p-ERK, and concomitant suppression of phospho-MYC (S62) and total MYC protein in the tumors within a week of Cobimetinib treatment initiation *in vivo* (**Fig. 4**). These findings, taken together with our phenotypic and signaling assessments (**Fig. 2** and **3**), strongly suggest that targeting the EphB2-MYC axis can be a rational therapeutic strategy against this aggressive malignancy.

**Figure 4.**
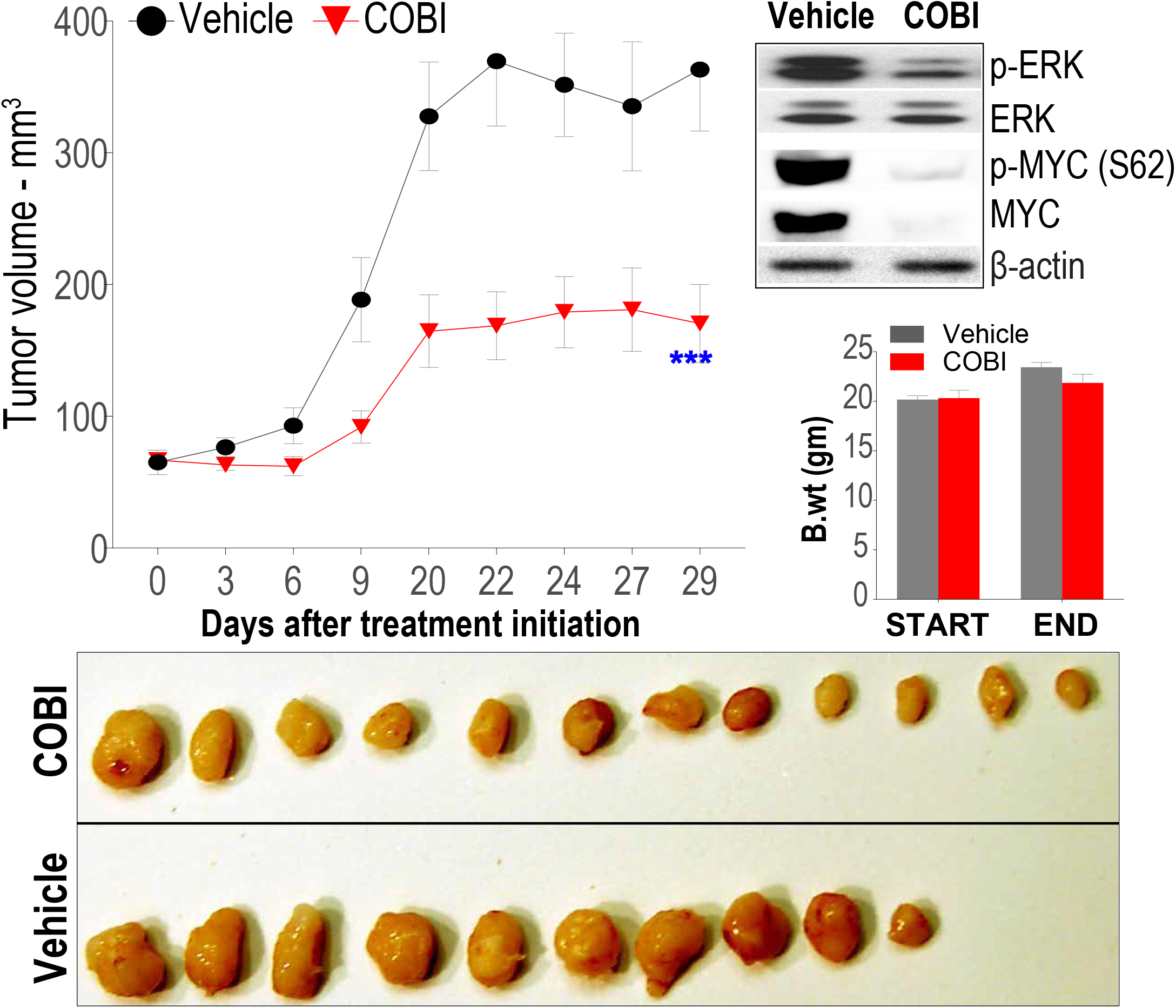
MEK inhibition suppresses MYC activity and EAC tumor growth *in vivo*. Assessment of tumor xenograft growth (left) of SKGT4 EAC cells in immune-deficient mice treated with COBI, as compared to vehicle control treatments. Y-axis depicts tumor volume in mm^3^ over time (X-axis) in EAC xenografts. Data are plotted as mean ± SEM estimated using at least 10 established xenograft tumors at day zero in respective arms per EAC cell line. *** (*P* < 0.0005) indicates significant differences in tumor volume at the final time-point between respective groups, estimated using a Student’s t-test assuming unequal variances. Western blot images (right) depicting protein levels of pharmacodynamic marker, p-ERK, and p-MYC/total MYC in representative tumor xenografts harvested after 1 week of treatment with either COBI or vehicle control. Note marked reduction of p-ERK, p-MYC, total MYC in COBI treated tumor xenografts as compared to vehicle. Bar graphs depict body weight assessments of COBI and vehicle treated mice at the beginning and at the end of the study. Photographic images (bottom) of harvested tumors from mice demonstrating the differences in tumor size in COBI versus vehicle treated mice at the final time-point.

## DISCUSSION

The alarming increase in the incidence of Barrett’s esophagus and esophageal adenocarcinoma (EAC) within the U.S., when coupled with the dismal survival rates for EAC on frontline chemoradiation therapy, and the limited avenues for targeted therapeutic interventions have exacerbated the need to decipher the molecular dependencies of this complex gastrointestinal disease. Here, we applied a systems biology framework to model signaling network activities using transcriptomic profiles of treatment-naïve primary patient samples, thus identifying hyperactivation of EphB2 signaling in both malignant as well as premalignant stages of the BE-EAC disease continuum, a finding that was further confirmed in a large independent cohort of patient samples (**Fig. 1**).

Notably, EphB2 is a member of the largest subgroup of receptor tyrosine kinases that are known to functionally impact diverse cellular processes during development and tissue homoeostasis (9). While the effects of Eph receptor activities in cancer are complex and paradoxical, with both decreased/increased expression (17) as well as higher mutational frequency (18) being linked to cancer progression across tissue contexts, this is the first report implicating hyperactivation of EphB2 signaling in the context of BE pathogenesis and its subsequent progression to EAC. Indeed, our observation of sustained activation of EphB2 signaling in primary patient lesions across the metaplasia-dysplasia-cancer stages of disease progression, when coupled with the striking growth suppression of BE/EAC cells upon *EPHB2* knockdown (**Fig. 2, Supplementary Fig. S1**), strongly support activation of EphB2 signaling as an early promoter of BE-EAC. Intriguingly, these findings are in stark contrast to the reported tumor-suppressive role of EphB2 signaling in other gastrointestinal malignancies such as colorectal cancer, where *EPHB2* is lost during the adenoma-carcinoma transition, and over-expression of *EPHB2* markedly suppresses colorectal cancer progression (19). These observations further underscore the distinct context-specific function of EphB2 signaling along the length of the gastrointestinal tract.

Our functional studies in BE/EAC cell line models (**Fig. 3A**) and quantitative assessments in primary BE and EAC lesions (**Fig. 1** and **3B**) further reveal MYC as a key mediator of EphB2 signaling. While EphB2 regulation of MYC has not been previously reported, prior studies indeed have implicated other Eph receptor family members as modulating MYC activity in diverse cancer contexts (20,21), intriguingly suggesting MYC as a critical downstream effector of Ephrin signaling in general. These findings have significant implications in the BE context given the likely role of MYC as an early promoter of BE pathogenesis. For example, a prior report in an organotypic culture model demonstrated that overexpression of MYC induces squamous to BE-like columnar transition (22), findings that are in line with our observation of enhanced MYC activity in transformative columnar cells in the porcine ESMG spheroid model (**Fig. 3C**). Moreover, deoxycholate, a key bile acid in gastroesophageal refluxate and a risk factor for BE, is shown to induce MYC protein expression (23). These findings suggest that cell-intrinsic/extrinsic factors that induce and/or sustain MYC activity likely play a determinant role in BE pathogenesis, and our study provocatively now connects this proto-oncogene to a potential master regulator, EphB2, in BE-EAC pathobiology.

While further studies are indeed warranted to fully decipher the precise intermediary steps in EphB2 regulation of MYC protein stability, our studies nonetheless offer new druggable targets for this aggressive cancer. In this regard, our preclinical studies with a clinically-approved MEK1 inhibitor, Cobimetinib, showed significant anti-tumor activity against EAC tumors, with concomitant inhibition of MYC activity, *in vivo* (**Fig. 4**). Moreover, given that Cobimetinib is already approved as an anti-cancer therapy in melanoma (24), our findings pave the way for rapid translational opportunities in EAC, a lethal cancer that has proven refractory to standard chemoradiation therapy. Since additional MEK inhibitors are in various phases of clinical development in other cancer contexts, our preclinical evaluations here provide a rationale for repurposing and evaluating MEK inhibitors in clinical trials as a potential targeted-therapeutic strategy in patients with EAC.

In summary, our investigations identify the EphB2-MYC signaling axis as a key molecular dependency in Barrett’s esophagus and esophageal adenocarcinoma, and reveal a novel therapeutic vulnerability in this increasingly-prevalent and aggressive esophageal malignancy.

## METHODS

Detailed methods are provided as Supplementary Methods.

### Patient samples

A discovery set of treatment-naïve 18 Non-Dysplastic Stable Barrett’s Metaplasia (NDSBM), 55 esophageal adenocarcinoma (EAC) and 11 paired normal esophageal squamous biopsy samples (nSQ) matching respective EACs was compiled for whole-transcriptome RNA sequencing (RNASeq) under an approved Institutional Review Board for Human Subjects Investigation protocol, as previously described (8). We similarly compiled an independent validation cohort consisting of 210 EAC, 133 Non-Dysplastic Barrett’s Metaplasia (NDBM), 57 Barrett’s Metaplasia with high-grade dysplasia (BE-HGD), 24 normal gastric (GAST) samples, and 461 normal squamous (nSQ) for qPCR-based validation studies.

### Estimation of signaling network activities in BE/EAC using the InFlo systems biology framework

Whole-transcriptome RNA sequencing (RNASeq) on the discovery cohort of EAC, NDSBM and nSQ samples was performed as previously described (8), generating an average of 150 million paired-end reads per sample. Differential activation of genome-scale signaling networks in NDSBM versus nSQ, as well as EAC versus nSQ was conducted using a unique systems biology framework (InFlo) that we recently developed (6,7). Briefly, the normalized, log2-transformed FPKM values across the 78 RNASeq samples in our discovery cohort was processed using InFlo by first comparing each individual sample against the control set of 11 nSQ samples. Signaling network components that were significantly activated (InFlo Activity Index > 0; Wilcoxon *P* ≤ 0.05) in >50% of both NDSBM as well as EAC as compared to nSQ samples were selected as being robust pathways activated in BE/EAC.

### Cell culture

Human EAC cell lines were cultured in either Dulbecco’s minimal essential medium (FLO-1) or Roswell Park Memorial Institute medium (SKGT4, EsoAd1, OE19, OE33 and Eso26) supplemented with 10% fetal bovine serum. Non-dysplastic (CP-A) and dysplastic (CP-B, CP-C, CP-D) human BE cell lines were cultured as previously described by us (7). Cell lines were tested for authenticity using STR genotyping, and were screened periodically for mycoplasma contamination.

### Generation of SKGT4 cells with stable knock-down of EPHB2

SKGT4 cells were infected with either the SMARTvector nontargeting or *EPHB2*-targeting lentiviral shRNA Tet-On 3G Doxycyline (Dox)-inducible system. SKGT4 cells were passaged 48 hours after infection and selected with 0.5 μg/mL puromycin for 2-3 weeks to obtain stable clones with nontargeting or *EPHB2* shRNA. Stable clones were treated with 1µg/ml doxycycline for 48-120 hours followed by assessment for efficient knockdown of EphB2 by immunoblotting.

### In vitro phenotypic assays

Cell growth assessments were quantified following seeding using the IncuCyte ZOOM automated live cell kinetic imaging system (Essen BioScience), as previously described (7). Colony growth assessments were quantified as previously described (7). Cell viability assessments were performed using the CellTiter-Glo 2.0 Cell Viability Assay. Significant differences in cell viability/growth/clonogenicity between test *versus* control groups were estimated using a Student’s t-test assuming unequal variances.

### InFlo-based identification of effector pathways of EphB2 signaling

EAC and dysplastic-BE cells were transfected for 48hrs with either siRNA directed against *EPHB2* or non-targeting/control siRNA, followed by genome-scale transcriptomic profiling. The resulting cell-line specific transcriptomic profiles were processed using InFlo by comparing the profiles of each of the *siEPHB2*-treated samples against the respective non-targeting controls. This resulted in the estimation of differential activities of signaling network components on a genome-scale, with negative values corresponding to lower activities and positive values corresponding to higher activities in *siEPHB2*-treated samples as compared to the respective non-targeting controls. Signaling network components that were significantly deregulated in the *siEPHB2*-treated samples across both EAC (SKGT4) and dysplastic-BE (CP-D) cells EAC and BE cells were subsequently selected to identify consensus downstream mediators of EphB2 signaling.

### In vivo tumor xenograft growth assay

Tumor xenograft growth assays were performed as previously described (7). Briefly, subcutaneous tumor xenografts were created by injecting either parental SKGT4 EAC cells or SKGT4 cells stably expressing either Dox-inducible *EPHB2*/Control shRNA constructs, suspended in 50% Geltrex, bilaterally (4×10^6^ cells per flank) into the flanks of 4–5-week-old female athymic Crl:NU(NCr)-Foxn1^nu^ mice. For shRNA xenograft studies, mice were switched to a 625 mg/kg doxycycline diet 24 hours after inoculation for the duration of the study. For MEK inhibitor studies, after the xenograft tumors reached a minimum size of 30mm^3^, the mice were randomized to be treated, via oral gavage, with either vehicle (0.5% Methylcellulose) or with vehicle containing 10 mg/kg of Cobimetinib, once daily. Tumor volumes were estimated 2-3 times weekly. A two-sided Student’s t-test, assuming unequal variances, was used to determine significant differences in tumor volumes across comparisons. All animal procedures were approved by the Case Western Reserve University Institutional Animal Care and Use Committee and followed NIH guidelines.

### Single-cell RNA sequencing analyses in a porcine 3D ESMG spheroid model

Porcine esophageal submucosal gland (ESMG) cultures were created as previously reported (14). For the single cell RNA sequencing, day 7 spheroids were collected from three pigs, representing a mix of two phenotypes: the solid squamous-like (SQ-like) spheroids and the hollow BE-like spheroids. A total of 2,677 single cells were recovered for single cell sequencing using the 10x Genomics Chromium System at the Duke Molecular Physiology Institute Molecular Genomics Core, followed by sequencing on an Illumina platform. The resulting paired-end reads were demultiplexed and aligned to the porcine genome reference (Sus Scrofa 11.1.101), followed by gene expression quantification and normalization resulting in a total of 16,279 porcine genes being detected in this scRNAseq run, with a median of 5,169 genes detected per cell. Graph-based clustering of the cells using the normalized single-cell gene expression profiles identified single-cell clusters, which were then evaluated for expression levels of predefined marker genes of normal squamous epithelium (TP63) and Barrett’s Metaplasia (KRT7), thus identifying two major single-cell cluster groups belonging to either the SQ-like (TP63^High^ & KRT7^Low^) or BE-like (KRT7^High^ & TP63^Low^) spheroids.

### Modeling EphB2 Signaling Activity in the porcine scRNASeq dataset

We recently extended our InFlo framework to be applicable to assess the activity levels of signaling networks using single-cell RNA sequencing data. The scRNASeq-based InFlo framework (scInFlo) is designed to take as input normalized single-cell gene expression profiles and estimate signaling network activities in individual cells as compared to a specific control population of cells. Accordingly, scInFlo was employed to determine EphB2 signaling activity in KRT7^High^ & TP63^Low^ (BE-like) cells as compared to KRT7^Low^ & TP63^High^ (SQ-like) cells in the porcine scRNASeq dataset described above.

### Assessment of MYC transcriptional activity

MYC activity levels in individual samples in the bulk RNASeq (human) datasets or individual cells in the porcine scRNASeq datasets was estimated using a statistical methodology designed to infer activity levels of transcription factors using relative expression levels of the transcription factor’s target genes (see Supplementary Methods). Of note, a total of 379 human MYC target genes or their porcine homologues were used to quantify MYC activity in individual samples/cells.

## ACKNOWLEDGEMENTS

We would like to thank Lakshmeswari Ravi and Aruna Kumar Chelluboyina for their technical input on the experimental methodologies employed in this study. We would like to thank City Packing in Burlington, NC, in particular Thomas McGarity and Jamie Corbett, for their generous contributions of porcine esophagus for this research. We also acknowledge the assistance of the Duke Molecular Physiology Institute Molecular Genomics Core at Duke University for the generation of single-cell RNA sequencing data included in this manuscript.

