## Supplementary Figures and Tables for "The EphB2-MYC Axis is a Major Determinant of Barrett’s Pathobiology and a Therapeutic Vulnerability in Esophageal Cancer"

**Fig. S1.** Impact of EphB2 on BE cell viability

**Fig. S2.** Signaling networks modulated upon siRNA-based knockdown of EPHB2 in BE/EAC cell lines

##### Supplementary Tables

**Table S1.** Sample distribution in the discovery RNASeq Cohort

**Table S2**. Sample distribution in the validation cohort

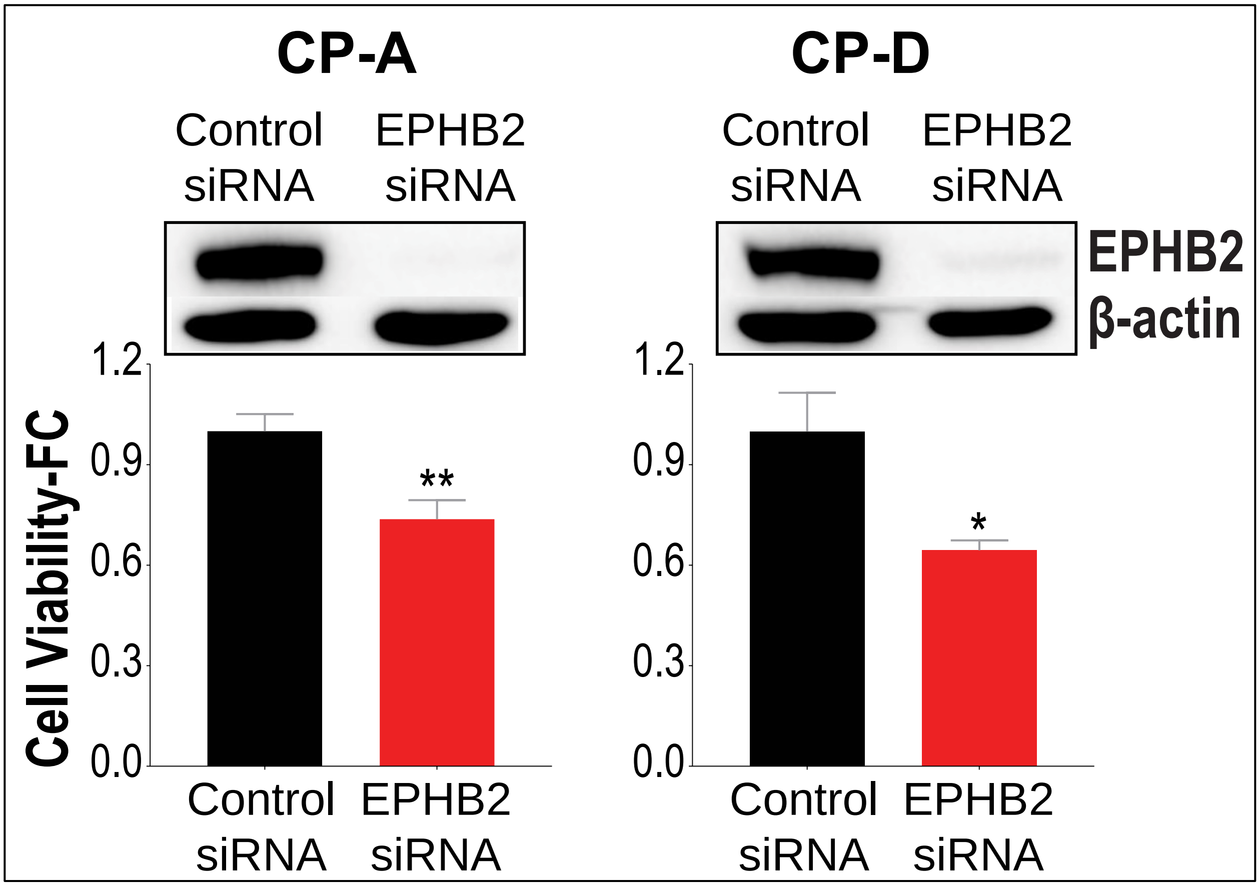


**Fig. S1**. **Impact of EphB2 on BE cell viability.** Bar graphs showing cell viability (Y-axis) upon siRNA-based EPHB2 knockdown in distinct metaplastic (CP-A) and dysplastic (CP-D) cell line models, expressed as a fraction of cell viability in cells treated with Control siRNA. Western blot images depict EPHB2 protein expression 48hrs following treatment with either EPHB2/Control siRNA. All data are plotted as mean ± SEM, obtained from three independent experiments. * ($P$ < 0.05) and ** ($P$ < 0.005) indicate significant differences in cell viability estimated using a Student's t-test assuming unequal variances.

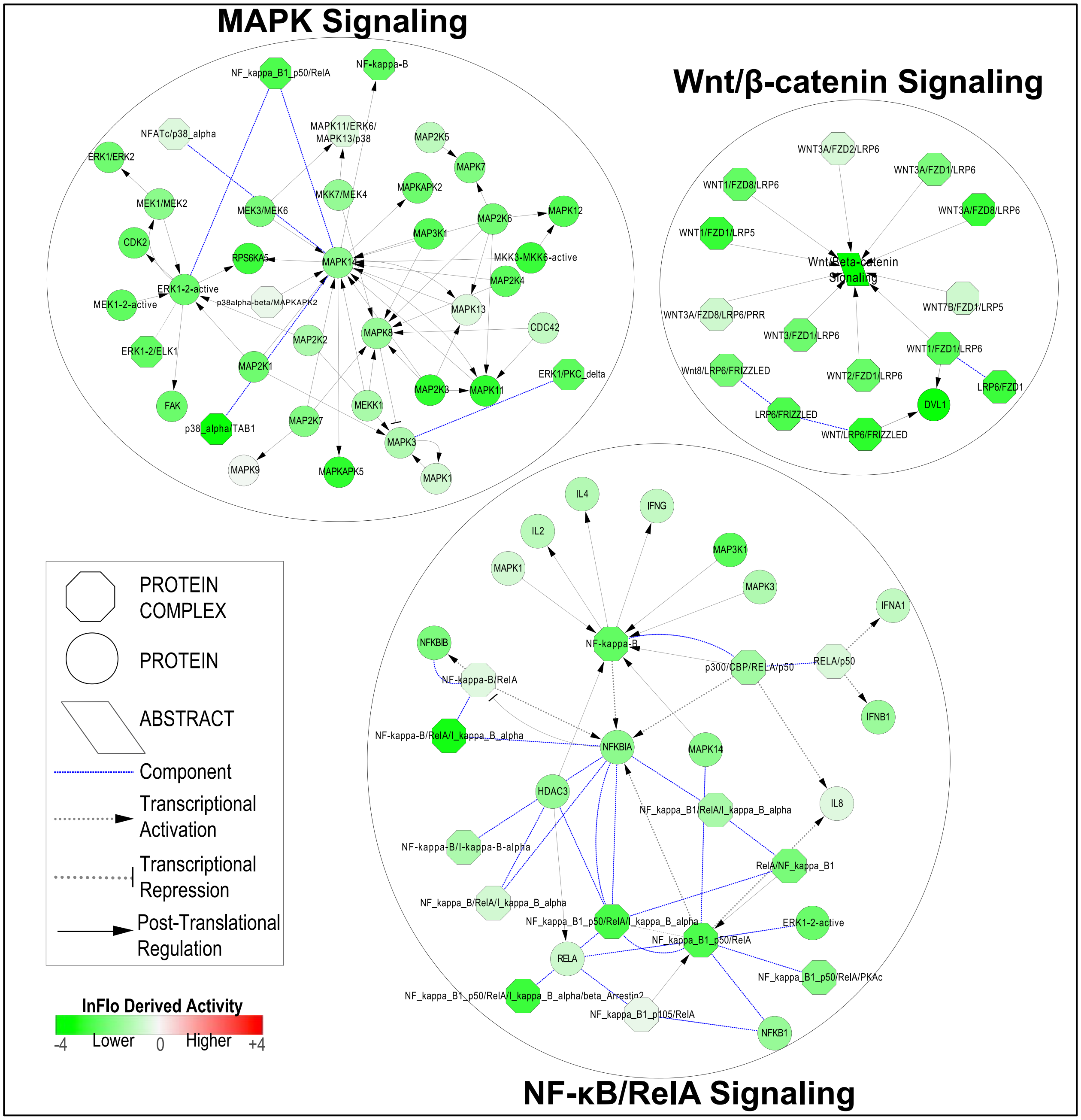


**Fig. S2. Signaling networks modulated upon siRNA-based knockdown of *EPHB2* in BE/EAC cell lines.** InFlo-based analysis of whole transcriptome RNASeq performed on EAC (SKGT4) and dysplastic-BE (CP-D) cells revealing marked reduction in the activities of signaling network components and their transcriptional targets in *EPHB2* siRNA treated cells as compared to non-targeting Control siRNA. Shown are the consensus signaling networks deregulated in both EAC and BE cells upon siRNA-based *EPHB2* knockdown. Note the marked reduction in the activities of NF-κB/RelA, Wnt/β-catenin and MAPK signaling components upon *EPHB2* KD in both EAC/BE cells.

**Table S1. Sample distribution in the discovery RNAseq Cohort**

| Tissue Type | Number of Samples | Median Age at Diagnosis (Range) | Gender Distribution | Cancer Stage Distribution |
|---|---|---|---|---|
| **Esophageal Adenocarcinoma (EAC) $** | 49 | 65 (36 - 88) | 89% (Male) 11% (Female) | Stage I (17.9%), Stage II (19.6%), Stage III (46.4%), Stage IV (16.1%) |
| **Non-dysplastic Stable Barrett's Metaplasia (NDSBM)*** | 18 | 56 (18-84) | 94% (Male) 6% (Female) | NA |
| **Normal Esophageal Squamous (nSQ) #** | 11 | 64 (45-83) | 90% (Male) 10% (Female) | NA |
| **Total** | **78** | | | |

$11% of EACs were gastroesophageal junctional adenocarcinomas
*Median surveillance of 9 years, ranging from 6 to 22 years
### Each of the 11 normal SQ samples was obtained from respective EAC patients included in the RNA sequencing

**Table S2. Sample distribution in the validation cohort**

| Tissue Type | Number of Samples | Median Age (Range) | Gender Distribution | Cancer Stage Distribution |
|---|---|---|---|---|
| **Esophageal Adenocarcinoma (EAC) ^** | 210 | 64 (34 - 89) | 77% (Male) 15% (Female) | Stage I (14.1%), Stage II (16.8%), Stage III (52.2%), Stage IV (15.0%) |
| **Normal Esophageal Squamous (nSQ)** | 461 | 64 (34 - 89) | 77% (Male) 15% (Female) | NA |
| **Non-Dysplastic Barrett's Metaplasia (NDBM) *** | 133 | 65.5 (36-93) | 71% (Male) 34% (Female) | NA |
| **BM with high-grade dysplasia (BE-HGD)** | 57 | 66 (46-80) | 89% (Male) 11% (Female) | NA |
| **Normal gastric (GAST)** | 24 | 63 (36-82) | 85% (Male) 15% (Female) | NA |
| **Total** | **885** | | | |

^13% of EACs were gastroesophageal junctional adenocarcinomas
*Clinical follow-up information unavailable (progression status unknown) for these patients
