## Supplementary Methods for "The EphB2-MYC Axis is a Major Determinant of Barrett’s Pathobiology and a Therapeutic Vulnerability in Esophageal Cancer"

### PATIENTS AND METHODS

#### *Patient samples*

We compiled a discovery set of treatment-naïve 18 Non-Dysplastic Stable Barrett's Metaplasia (NDSBM), 55 esophageal adenocarcinoma (EAC) and 11 paired normal esophageal squamous biopsy samples (nSQ) matching respective EACs for whole-transcriptome RNA sequencing (RNASeq). Study subjects were recruited under an approved Institutional Review Board for Human Subject Investigation protocol across institutions participating in the Barrett's Esophagus Translational Research Network (BETRNet). All tissues were collected from patients with Barrett's Metaplasia (BM) or newly diagnosed EAC undergoing endoscopy prior to receiving therapy, with endoscopic biopsies being snap frozen at bedside and stored at -80°C to preserve RNA. NDSBM biopsies were obtained from subjects with at least 2cm of BM who had no history of dysplasia and have not developed dysplasia or cancer upon subsequent follow-up (median surveillance of 9 years, ranging from 6 to 22 years). Biopsies from NDSBM patients were obtained preferentially from areas most likely to have intestinal metaplasia using high definition white light and narrow band imaging guidance (1-3). EAC biopsies were obtained preferentially from areas of visible cancer that did not appear ulcerated or necrotic at endoscopy. nSQ biopsies, matching the EAC cases, were obtained a minimum of 3cm proximal to the squamo-columnar junction. In each case, sister biopsies, obtained adjacent to the frozen biopsies, were formalin-fixed and paraffin-embedded for subsequent histological examination by an expert anatomic pathologist (J.E.W.) to confirm the diagnosis. Histopathological evaluation of formalin-fixed paraffin-embedded biopsy sections showed a cancer cell content of greater than or equal to 50% in EACs (J.E.W.).

We similarly compiled an independent validation cohort from the participating BETRNet institutions consisting of 210 EAC, 133 Non-Dysplastic Barrett's Metaplasia (NDBM), 57 Barrett's Metaplasia with high-grade dysplasia (BE-HGD), 24 normal gastric (GAST) samples, and 461 normal squamous (nSQ). Of note, the majority of nSQ biopsies were patient-matched to the EAC/NDBM/HGD samples and the normal gastric biopsies were obtained from the gastric fundus in patients with NDBM/EAC included in the validation cohort.

Patient demographics including age, gender and cancer stage distribution for both the discovery and validation cohorts are included in Supplementary Tables S1 and S2. As outlined in Supplementary Tables S1 and S2, women subjects were included in the study, but since BE and EAC largely occur in Caucasian males, the majority of patients in our cohorts were Caucasian males.

***Whole-transcriptome RNA sequencing and transcript abundance quantification***

Whole-transcriptome RNA sequencing (RNASeq) on the discovery cohort of EAC, NDSBM and nSQ samples was performed as previously described by us (4). Briefly, total RNA was extracted using the *mir*Vana kit (Ambion, Austin, TX), and RNA concentration was measured using a NanoDrop 2000C spectrophotometer (Thermo Scientific, Wilmington, DE) and samples checked for integrity using an Agilent BioAnalyzer 2100 (Agilent Technologies, Santa Clara, CA) prior to sequencing. The minimum RNA integrity number (RIN) value was 6.0 for the RNASeq samples with an average RIN value of 8.0. RNASeq was performed by the Oklahoma Medical Research Foundation Next Generation Sequencing Core Facility (Oklahoma City, OK). RNASeq library preparations were performed using the Illumina Truseq RNA sample preparation kit as per the manufacturer's protocol (Illumina, Inc., San Diego, CA). Briefly, poly-A RNA was enriched from total RNA via hybridization with biotinylated poly-dT oligomers. The enriched poly-A RNA was converted into cDNA and sheared to an average of 300bp using a Covaris S2 sonicator (Covaris, Inc., Woburn, MA). Sonicated cDNA was end repaired, single-base A-tailed, ligated with Illumina-specific adapters, and amplified with PCR to enrich for successful ligation products. Libraries were pooled in sets of 6 prior to absolute quantification by quantitative real-time PCR (qPCR) on a Roche Lightcycler 480 (F. Hoffmann-La Roche AG, Basel, Switzerland) using the Kapa Biosystems Library Quantification kit (Kapa Biosystems, Inc., Wilmington, MA). Library pools were sequenced across 3 lanes of an Illumina Hiseq 2500 instrument (Illumina, Inc.) using paired-end 100bp reads, generating an average 150 million reads per sample. Sequence data were mapped using STAR (ver 2.7.0a) (5,6) and transcript abundance in FPKM units (Fragments per kilo-base of mRNA per million reads) estimated for a total of 22,160 unique coding genes and 34,449 unique transcripts from RefSeq using Cufflinks (ver 2.2.1) (7).

***Estimation of signaling network activities in BE/EAC using the InFlo systems biology framework***

Additionally, differential activation of genome-scale signaling networks in NDSBM versus nSQ, as well as EAC versus nSQ were conducted using a unique systems biology framework (InFlo) that we recently developed. InFlo (8,9) can robustly integrate diverse high-throughput molecular measurements to decode mechanisms contributing to complex disease phenotypes. InFlo employs a unique probabilistic machine-learning framework that leverages sparse molecular measurements to infer signaling network activities on a genome scale that are not explicitly measured by routine molecular profiling platforms. We used InFlo to identify pathways that exhibit differential activities in NDSBM or EAC samples by first comprehensively characterizing global pathway network activities across each of the NDSBM, EAC, and nSQ samples in our discovery cohort. Briefly, gene expression profiles across the 78 RNASeq samples in our discovery cohort was processed using InFlo by first comparing each individual sample against the control set of 11 nSQ samples. This resulted in the estimation of a differential activity level for each component of the pathway networks, with negative values corresponding to lower activity and positive values corresponding to higher pathway-specific activity as compared to the nSQ control set of samples. In order to identify signaling networks deregulated in NDSBM or EAC, we first identified sub-networks that were significantly deregulated in NDSBM versus nSQ, and EAC versus nSQ, respectively, using the non-parametric Wilcoxon rank-sum test. Pathway network components with a non-parametric Wilcoxon p-value $\leq 0.05$ were considered significant. Network components that were significantly activated (InFlo Activity Index > 0) in >50% of both NDSBM as well as EAC samples were selected as being robust pathways activated in BE/EAC. Strikingly, our InFlo-based analysis recapitulated many of the known molecular pathways involved in BE/EAC-pathogenesis, including activation of MYC signaling (10,11), CDX1/CDX2 signaling (11-15), NF-κB/RelA signaling (16,17) and Trefoil Factor 3 (TFF3) signaling (18) as being activated in the majority of NDSBMs as well as EACs. In addition to these well-known pathway networks, InFlo additionally identified EphB2 signaling as being hyperactivated (InFlo Activity Index > +2) as compared to nSQ in 100% of NDSBM and 88% of EACs.

***Quantitative real-time PCR***

1 µg of total RNA was reverse-transcribed using Superscript III First-Strand Synthesis (Life Technologies, Grand Island, NY, #18080) according to the manufacturer's protocol. The cDNA templates were diluted 1:20 with Ultrapure distilled water (Life Technologies, #10977) for a final concentration of 1.25 ng/µL. qPCR analysis was performed using iQ SYBR Green Supermix system (Bio-Rad Laboratories, Hercules, CA, #170-8887) with pre-designed intron-spanning primer sets for *EPHB2 (*Qiagen Inc., Valencia, CA, #QT00089495) and *cMYC* (Bio-Rad Laboratories, Hercules, CA, Biorad, #10041595). *B2M* (Qiagen, #QT00088935) was used as an endogenous qPCR control for all samples. qPCR reactions were carried out in 25 µL volume containing 2.5 ng of cDNA template, with the following PCR conditions; 95$^{o}$C for 5 minutes, and 40 cycles of 95$^{o}$C for 15 seconds, 55 $^{o}$C for 20 seconds, and 72 $^{o}$C for 20 seconds, followed by a melting curve analysis using a Bio-Rad CFX96 Real-Time PCR machine (Bio-Rad). Each qPCR reaction was carried out in triplicate and the mean expression levels across the triplicates were normalized to the mean expression values of *B2M* using the ΔCt method (mean test mRNA Ct (minus) mean *B2M* mRNA Ct). The log2 fold changes of *EPHB2* expression in NDBM (*N*=133), BE-HGD (*N*=57), EAC (*N*=210) and GAST (*N*=24) were expressed relative to the median expression of *EPHB2* in the SQ (*N*=461) samples. Of note, all of the above qPCR reactions and analyses were performed in accordance with the MIQE guidelines as previously described (19). All qPCR reactions were also performed in a blinded fashion to sample identity. A two-sided Student's t-test, assuming unequal variances, was used to determine significant differences in *EPHB2* expression between NDBM, HGD, EAC or GAST versus SQ samples. The sample-size of the validation cohort provided >80% power to detect at least a 2-fold difference in *EPHB2* expression between NDBM, HGD, EAC versus SQ samples.

#### *Immunohistochemistry analysis on primary tissue samples*

Briefly, slides containing 5µm formalin-fixed paraffin-embedded tissue sections from representative normal SQ, NDBM, BE-HGD and EAC cases were baked for 75 minutes at 60$^{0}$C and de-paraffinized in xylene, passed through graded ethanols and rehydrated in distilled water. Heat-induced antigen retrieval was carried out in a pressure cooker in citrate solution (pH 6.0) at 123$^{0}$C for 30 seconds and endogenous peroxidase was quenched with 3% hydrogen peroxide for 8 minutes. Charged proteins and endogenous IgG was blocked in background sniper solution for 20 minutes. Slides were incubated with the primary EphB2 antibody (1:200 dilution, Cell Signaling Technology Inc., Danvers, MA, # 83029) overnight at 4$^{0}$C. Detection of antibodies was achieved by incubating the sections with goat-anti rabbit horseradish peroxidase (HRP) conjugated polymers for 30 minutes. Sections were stained with 3,3'-Diaminobenzidine chromogen for 5 minutes to visualize antibody-antigen binding, and hematoxylin was used as a nuclear counterstain. In parallel, as a negative control, serial sections were similarly processed as above with the exclusion of primary antibodies.

#### *Cell Culture*

Human esophageal adenocarcinoma (EAC) cell line FLO-I (Millipore Sigma, St Louis, MO #11012001) was cultured in Dulbecco's minimal essential medium (DMEM, Life Technologies, #11965) supplemented with 10% fetal bovine serum (FBS, Hyclone Laboratories, Logan, UT, #SH30071). Human EAC cell lines JH-EsoAd1 (kindly provided by Drs. James Eshleman, Johns Hopkins Medical Institutions, Baltimore, MD and Anirban Maitra, MD Anderson Cancer Center, Houston, TX), ESO26, SKGT4, OE19 and OE33 (Millipore Sigma, #11012009, #11012007, #96071721, #96070808) were cultured in Roswell Park Memorial Institute medium (RPMI, Life

Technologies, #1875-119) supplemented with 10% FBS. Normal squamous esophageal cell line (EPC2, kindly provided by Dr. Anil Rustgi, Columbia University, New York, NY), non-dysplastic (CP-A) and dysplastic (CP-B, CP-C, CP-C) human BE cell lines (American Type Culture Collection, Manassas, VA, #CRL-4027, #CRL-4028, #CRL-4029, #CRL-4030) were cultured as previously described (20). Cell lines were tested for authenticity using short tandem repeat (STR) genotyping prior to experimental use. Cell lines were also screened periodically for mycoplasma contamination using the MycoAlert kit (Lonza, Basel, Switzerland, # LT07-518).

***Generation of SKGT4 cells with stable knock-down of EPHB2***

SKGT4 cells were infected with either the SMARTvector nontargeting or *EPHB2*-targeting lentiviral shRNA Tet-On 3G Doxycyline (Dox)-inducible system (Dharmacon Inc, Lafayette, CO, #VSC6571 and #V3SH7675-08EG2048). Briefly, SKGT4 cells were passaged 48 hours after infection and selected with 0.5 µg/mL puromycin (Life technologies, #A1113803) for 2-3 weeks to obtain stable clones with nontargeting or *EPHB2* shRNA. Clones selected from 6 individual *EPHB2* shRNA were treated with 1µg/ml doxycycline (Takara Bio, Mountain View, CA, #631311) for 48-120 hours and assessed independently for efficient knockdown of EphB2 protein by western blotting. Of the 6 *EPHB2* shRNAs tested, only one shRNA exhibited strong EphB2 knockdown and was thus selected for subsequent *in vitro* and *in vivo* assays.

***Protein extraction and Immunoblotting***

Cells were seeded at a density of $5x10^4$ cells/well – $2.5x10^5$ cells/well in a 6-well plate in serum-free or 2% FBS-supplemented RPMI media. Total protein was extracted 6 hours after treatment with 10µM Cobimetinib (MedChemExpress, #HY-13064A) or 24-48 hours after treatment with

50nM siRNAs, using cold cell lysis buffer (Cell Signaling Technology, Danvers, MA, #9803) supplemented with protease (Millipore Sigma, #04693124001) and phosphatase inhibitor cocktail (Millipore Sigma, #04906845001). Lysates were then centrifuged at 15,000g for 15 minutes at 4˚C and the supernatant was used for immunoblotting. Protein concentration was quantified using BCA Protein Assay Kit (Life Technologies, #23225). 10 to 30 μg of protein was loaded into one lane of a 4-12% polyacrylamide gel (Life Technologies, #NP0321BOX) and subjected to sodium dodecyl sulfate–polyacrylamide gel electrophoresis and transferred to polyvinylidene difluoride membrane (Biorad Laboratories, #1704156). Membranes were blocked with 5% milk or 5% bovine serum albumin (The Jackson Laboratory, Bar Harbor, ME, #001-000-161) in TTBS (0.05% Tween-20 in tris buffered saline) and incubated with primary antibody overnight at 4ºC at a dilution of 1:1000 in 5% BSA in TTBS for EphB2 (Cell Signaling Technology, # 83029), EFNB1/B2 (Invitrogen, # PA5-105732), Phospho-c-MYC (Ser62) (Cell Signaling Technology, # 13748), total c-MYC (Cell Signaling Technology, #5605), Phospho-p44/42 MAPK (Erk1/2) (Thr202/Tyr204) (Cell Signaling Technology, #9101), total p44/42 MAPK (Erk1/2) (Cell Signaling Technology, #9102). Blots were incubated with anti-rabbit HRP secondary antibodies (Cell Signaling Technology, #7074) at 1:5000 in 5% milk in TTBS. Membranes were stripped using Restore Plus (Thermo Scientific, Waltham, MA, #46430) and re-probed with mouse anti-β-actin (Cell Signaling Technology, #3700) and anti-mouse HRP (Cell Signaling Technology, #7076) at 1:10,000 as a loading control. Chemiluminescence was visualized using ECL Western Blotting Detection Kit (Global Life Sciences Solutions USA LLC, Marlborough, MA, #RPN2232, #RPN2235) and quantified using the Image Lab software (Bio-Rad Laboratories, Hercules, CA, # 12012931).

#### *Cell proliferation assays*

Cell proliferation assessments were quantified as previously described by our group (9). Cells were seeded at a density of 2000-5000 cells per well in a 96-well plate. After 24 hours, cells were transfected with 50nM siRNA (ON-TARGETplus SMARTpool) directed against *EPHB2* (Dharmacon Inc., #L-003122), *MYC* (Dharmacon Inc., # L-003282) or non-targeting/control siRNA (Dharmacon Inc., #D-001810), using RNAiMAX transfection reagent transfection reagent (Life Technologies, #13778075) in RPMI supplemented with 2% or 10% fetal bovine serum. Growth measurements expressed as percent confluency were obtained over the course of 120 hours following seeding using the IncuCyte ZOOM automated live cell kinetic imaging system (Essen Instruments, Ann Arbor, MI). Significant differences in cell growth between respective treatment versus control groups were estimated using a two-sided Student's t-test assuming unequal variances.

#### *Cell viability assay*

Cells were seeded at a density of 2000-5000 cells per well in a 96-well plate. After 24 hours, cells were transfected with 50nM siRNA (ON-TARGETplus SMARTpool) directed against *EPHB2* (Dharmacon Inc., #L-003122), *MYC* (Dharmacon Inc., # L-003282) or non-targeting/control siRNA (Dharmacon Inc., #D-001810), using RNAiMAX transfection reagent transfection reagent (Life Technologies, #13778075) in RPMI supplemented with 2% or 10% fetal bovine serum. At the end of 120 hours, cells were assessed for ATP activity using CellTiter-Glo 2.0 Cell Viability Assay (Promega, Madison, WI, #G9241). A two-sided Student's t-test, assuming unequal variances, was used to determine significant differences in cell viability across comparisons.

#### *Clonogenic assays*

Colony growth assessments were quantified as previously described by our group (9). Cells were seeded at densities of 250-2500 cells per well in a 6-well plate, respectively, and maintained in culture overnight in RPMI supplemented with 10% FBS. After 24 hours, cells were transfected with 50nM siRNA (ON-TARGETplus SMARTpool) directed against *EPHB2* (Dharmacon Inc., #L-003122) or nontargeting/Control siRNA (Dharmacon Inc., #D-001810), using RNAiMAX transfection reagent (Life Technologies, #13778075). The siRNA treatments were terminated after 48 hours, following which the cells were maintained in RPMI media supplemented with 10% FBS throughout the duration of the experiment. For the shRNA-based *EPHB2* clonogenic assay, SKGT4 cells stably expressing either Dox-inducible *EPHB2*-shRNA or Dox-inducible Control-shRNA were plated at densities of 300-500 cells per well in a 6 well plate in RPMI media supplemented with 10% FBS. After 24 hours, cells were treated with 1µg/µl doxycycline; following which they were maintained in dox-containing media throughout the duration of the experiment. 7-14 days after initial seeding, colonies were stained as previously described (21) and quantified using Alpha imager (Alpha Innotech, San Leandro, CA). A two-sided Student's t-test, assuming unequal variances, was used to determine significant differences in colony numbers across comparisons.

#### *In vivo tumor xenograft growth assay*

Tumor xenograft growth assays were performed as previously described (21). Briefly, subcutaneous tumor xenografts were created by injecting either parental SKGT4 EAC cells, or SKGT4 cells stably expressing either Dox-inducible *EPHB2*/Control shRNA constructs, suspended in 50% Geltrex (Life Technologies, #A1413201) into the bilateral flanks ($4x10^6$ cells per flank) of 4–5-week-old female athymic Crl:NU(NCr)-Foxn1$^{nu}$ mice (Charles River, Wilmington, MA). 5-10 mice were included per experimental or control arm. For shRNA studies, the next day after inoculations, mice feed were changed to 625 mg/kg doxycycline diet (Envigo, Indianapolis, IN, #TD.01306) and maintained on this diet for the duration of the study. Tumor volumes were estimated using the formula [1/2 (length x width$^2$)] 2-3 times weekly. For MEK inhibitor studies, after the xenograft tumors reached a minimum size of 30mm$^3$, the mice were randomized to be treated, via oral gavage, with either vehicle (0.5% Methylcellulose, Millipore Sigma, #64632) or with vehicle containing 10 mg/kg of Cobimetinib (MedChemExpress, Princeton, NJ, #HY-13064A), once daily. Representative tumors from each treatment arm were harvested after one week of treatment followed by protein assessments by immunoblotting as outlined above. A two-sided Student's t-test, assuming unequal variances, was used to determine significant differences in tumor volumes across comparisons. All animal procedures were approved by the Case Western Reserve University Institutional Animal Care and Use Committee and followed NIH guidelines.

***InFlo-based identification of effector pathways of EphB2 signaling***

EAC (SKGT4) and dysplastic-BE (CP-D) cells were seeded at a density of 100,000 cells per well in 6-well plates. After 24 hours, cells were transfected with 50nM siRNA (ON-TARGETplus SMARTpool) directed against *EPHB2* (Dharmacon Inc., #L-003122) or non-targeting/control siRNA (Dharmacon Inc., #D-001810), using RNAiMAX transfection reagent transfection reagent (Life Technologies, #13778075) in RPMI supplemented with 2% fetal bovine serum (for SKGT4) or keratinocyte media (for CP-D). siRNA treatments were performed, in triplicate, for SKGT4 and CP-D cell line models, followed by RNA extraction 48hrs post siRNA treatment and RNA-sequencing, resulting in an average of 60 million paired-end reads per replicate across cell line models and siRNA treatments. The resulting gene expression profiles were processed using InFlo by first comparing the transcriptome profiles of each of the *siEPHB2*-treated samples against the 3 non-targeting siRNA samples for SKGT4 and CP-D cell lines, respectively. This resulted in the estimation of differential activities of signaling network components on a genome-scale, with negative values corresponding to lower activities and positive values corresponding to higher activities in siEPHB2-treated samples as compared to the respective non-targeting controls. Signaling network components that were consistently and markedly down-/up-regulated in the siEPHB2-treated samples across both EAC (SKGT4) and dysplastic-BE (CP-D) cells were selected to identify consensus downstream mediators of EphB2 signaling.

***VIPER-based assessments of MYC activity using transcriptomic profiling of MYC target genes***

MYC activity levels in individual bulk RNASeq samples were estimated using the VIPER statistical methodology (22). VIPER is designed to infer activity levels of transcription factors using relative expression levels of the transcription factor's target genes as curated in the DoRothEA regulon database (23). Similarly, MYC activity in individual cells within the porcine 3D esophageal submucosal gland single-cell RNA sequencing dataset (see below) were estimated as outlined in the recent benchmarking study on transcription factor activity assessments in single-cell RNAseq datasets (24). Of note, a total of 379 porcine genes were identified as homologues of the human MYC target genes listed in the DoRothEA regulon database and used to quantify MYC activity in individual cells in the porcine scRNASeq dataset.

***Analysis of a single-cell RNA sequencing (scRNASeq) dataset of a porcine 3D ESMG spheroid model***

Porcine esophageal submucosal gland (ESMG) cultures were created as previously reported (25). In brief, under a North Carolina Dept. of Agriculture permit, research specimens of porcine esophagus were obtained from City Packing in Burlington, NC. Samples were kept cold in PBS during transport to the lab, where ESMGs were carefully dissected, leaving the squamous mucosal intact. After dissection into cold PBS, ESMGs were minced and then incubated in minimal media with antibiotics: Advanced Dulbecco's modified Eagle medium/F12 (Thermo Fisher) with metronidozole 10 mg/mL (Sigma), Primocin (InvivoGen), HEPES 10 mmol/L (Thermo Fisher), and Glutamax 1× (Thermo Fisher). After 15 minutes at 37°C, dithiothreitol was added (to 3 mmol/L) for an additional 10 minutes. Following incubation, the mixture of minced glands was centrifuged at 100 × g and rinsed with minimal media, then resuspended in 480 μL Growth Factor Reduced Matrigel Matrix (Corning, Durham, NC) (GFR Mtatrigel) in order to create 20-μL patties of ESMGs in Matrigel for 3D culture. After plating, the Matrigel patties were overlaid with 800 μL minimal media that was changed every 2–3 days.

After seven days in culture, Matrigel patties were mechanically loosened, collected, and chilled for 10 min over ice in cold minimal media. This was centrifuged (5 min at 1000 rmp) and incubated with trypsin (6mL per combined Matrigel patties from 24-well plate) for 15 minutes at room temperature with vortexing to disrupt the Matrigel. Cells were mixed by pipetting within a 15mL conical tube then centrifuged at 1000rpm for five minutes and resuspended in 12mL minimal media. The resuspended cells were filtered through a 70micrometer filter and then centrifuged at 100 rpm for five minutes, then counted in a hemocytometer before being plated at a density of 10,000 cells per 10 microliters of GFR Matrigel. After Matrigel patties solidified, 250 microliters of spheroid media (DMEM/F12 Advanced, 10mM HEPES, 1X Primocin, 1X Glutamax, N2, B27 and EGF) was added to each well of a 48 well plate. Again, media was changed every 2-3 days.

For the single cell RNA sequencing, day 7 spheroids were collected from three pigs, representing a mix of two phenotypes the solid squamous-like spheroids and the hollow BE-like spheroids. Single cells were dissociated from spheroids, washed and resuspended in 1x PBS / 0.04% BSA solution at a concentration of 1000 cells per microliter. A target of 5,000 pooled cells was used and after size selection (50nm) and washing, 2,677 cells were recovered for single cell sequencing using the 10x Genomics Chromium System at the Duke Molecular Physiology Institute Molecular Genomics Core. A Cellometer (Nexcelom) used to determine the cell viability and concentration. Cells were then combined with a master mix containing reverse transcription reagents. Gel beads carrying the Illumina TruSeq Read 1 sequencing primer, a 16bp 10x barcode, a 12bp unique molecular identifier (UMI) and a poly-dT primer were loaded onto the chip with oil for the emulsion reaction. The Chromium Controller partitioned cells into nanoliter-scale gel beads in emulsion (GEMS) for the reverse-transcription occurs. All cDNAs from one cell, shared a common barcode. After the RT reaction, full length cDNAs were cleaned with both Silane Dynabeads and SPRI beads. After purification, the cDNAs were assayed on an Agilent 4200 TapeStation High Sensitivity D5000 ScreenTape for qualitative and quantitative analysis. Enzymatic fragmentation and size selection were used to optimize the cDNA amplicon size. Illumina P5 and P7 sequences, a sample index, and TruSeq read 2 primer sequence were added via End Repari, A-tailing, Adaptor Ligation, and PCR. The final libraries contained P5 and P7 primers used in Illumina bridge amplification. Sequence was generated using paired end sequencing (one end to generate cell specific, barcoded sequence and the other to generate sequence of the expressed poly-A tailed mRNA) on an Illumina sequencing platform at a minimum of 50k reads/cell.

This resulted in a total of 217,931,948 paired-reads (Read1 length 28bp; Read2 length 91bp). These reads were subsequently demultiplexed, aligned and analyzed for differential activation of MYC signaling in single-cells derived from solid squamous-like (SQ-like) versus hollow Barrett's Esophagus-like (BE-like) spheroids (25,26) as outlined next. First, the raw sequence data were mapped to the porcine genome reference (Sus Scrofa 11.1.101), followed by barcode and UMI metrics using the CellRanger (3.1.0) pipeline (27). The CellRanger analysis of UMI counts versus Barcodes estimated that 92.5% of all reads originated from a total of 2,677 single cells with 81,409 mean reads and 28,541 median UMI counts per cell. Notably, a total of 16,279 porcine genes were detected in this scRNAseq run, with a median of 5,169 genes detected per cell. Cells with high percent of mitochondrial genes (>5%) were considered to be non-viable and excluded from all downstream analyses. Subsequently, the 10x data matrixes were imported into Seurat V3.0 R package (https://satijalab.org/seurat) to perform data filtration, gene expression quantification, normalization, dimension reduction and data visualization. Briefly, the scTransform framework for normalization and variance stabilization was employed for gene expression quantification and normalization, thus mitigating potential technical artifacts such as correlations between gene expression levels and total sequencing depth of a cell. The resulting normalized single-cell gene expression profiles were analyzed using Seurat's dimensionality reduction "RunPCA" function, followed by graph-based clustering of the cells using top 10 principle components that captured most of the variation in the data. The resulting single-cell clusters were subsequently visualized using Uniform Manifold Approximation and Projection for Dimension Reduction (UMAP) algorithm. The resulting single-cell clusters were then evaluated for expression levels of predefined marker genes of normal squamous epithelium (TP63) and Barrett's Metaplasia (KRT7), thus identifying two major single-cell cluster groups belonging to SQ-like ($TP63^{High}$ & $KRT7^{Low}$) or BE-like ($KRT7^{High}$ & $TP63^{Low}$) spheroids (25,26). Given the potential confounding effects of cell cycle heterogeneity on differential expression assessments, cell cycle phase scores were assigned to each cell based on the mean expression levels of 43 "S phase" and 54 "G2/M phase" marker genes (28) using the *CellCycleScoring* function in the *Seurat* package and cells expressing neither G2/M or S phase markers were deemed not cycling, thus resulting in a predicted classification of each cell into either G2M, S or G1 phase.

***Modeling EphB2 Signaling Activity in the porcine scRNASeq dataset***

We recently extended our InFlo framework to be applicable to assess the activity levels of signaling networks using single-cell RNA sequencing data (manuscript under preparation). The scRNASeq-based InFlo framework (scInFlo) is designed to take as input scTransform normalized single-cell gene expression profiles and estimate signaling network activities in individual cells as compared to a specific control population of cells. Accordingly, scInFlo was employed to determine EphB2 signaling activity in $KRT7^{High}$ & $TP63^{Low}$ (BE-like) cells as compared to $KRT7^{Low}$ & $TP63^{High}$ (SQ-like) cells in the porcine 3D ESMG model scRNASeq dataset described above. Briefly, scInFlo approximated a non-linear manifold defined by the transcriptomic profiles of SQ-like cells using a multi-kernel framework. Specifically, the normalized gene expression vectors of BE-like cells and SQ-like cells were mapped into a high-dimensional kernel space using multiple Gaussian kernels, $i.e., k(x,y) = \exp\left(-\frac{\|x-y\|^2}{\sigma}\right); x, y$ are BE/SQ-like cells, with each kernel defined by a unique $\sigma$ value ranging from 0.01 to 3, thus enabling the projection of each BE-like cell on to a minimum of 100 dimensions of Kernel Principal Component (KPC) spaces defined by the SQ-like cells. The resulting non-linear SQ-like cellular components of BE-like cells in the kernel space were inverse-mapped to obtain gene expression profiles for each BE-like cell, thus allowing for robust genome-scale differential expression assessments of each BE-like cells as compared to the corresponding nearest-neighbor SQ-like cellular components ($N$=15), with statistical significance probabilities being estimated using a Wilcoxon rank sum test. Given the sparsity of single-cell gene expression profiles, the probabilities of differential expression across genes for a given BE-like cell were adjusted using respective fold-change estimates corresponding to the nearest SQ-like cellular components using a soft function $\frac{1}{1+\exp\left(\frac{fc\hat{}2}{0.01}\right)}; where\ fc$ is the estimated log2 fold-change. The resulting genome-scale normalized differential expression assessments were combined with regulatory relationships derived from pathway network databases to infer the activities of EphB2 signaling in individual BE-like cells by combining differential expression assessments of individual components within the EphB2 signaling network as previously outlined within the core InFlo methodology (8).
